# Asymmetric genome merging leads to gene expression novelty through nucleo-cytoplasmic disruptions and transcriptomic shock in *Chlamydomonas* triploids

**DOI:** 10.1101/2024.08.06.604315

**Authors:** Lucas Prost-Boxoen, Quinten Bafort, Antoine Van de Vloet, Fabricio Almeida-Silva, Yunn Thet Paing, Griet Casteleyn, Sofie D’hondt, Olivier De Clerck, Yves Van de Peer

## Abstract

Genome merging is a common phenomenon in many organisms, causing a wide range of consequences on phenotype, adaptation, and gene expression, among other effects, yet its broader implications are not well understood. Two consequences of genome merging on gene expression remain poorly understood: dosage effects and evolution of expression. In this study, we employed *Chlamydomonas reinhardtii* as a model to investigate the effects of asymmetric genome merging by crossing a diploid with a haploid strain to create a novel triploid line. Five independent clonal lineages derived from this triploid line were evolved for 425 asexual generations in a laboratory natural selection (LNS) experiment. Utilizing fitness assays, qPCR, and RNA-Seq, we assessed the immediate consequences of genome merging and subsequent evolution over time. Our findings reveal substantial alterations in gene expression, protein homeostasis (proteostasis) and cytonuclear stoichiometry. Notably, gene expression exhibited expression level dominance and transgressivity (*i.e.*, expression level higher or lower than either parent). Ongoing expression level dominance and a pattern of “functional dominance” from the haploid parent was observed, alongside remarkable stability in expression patterns across generations. Despite major nucleo-cytoplasmic disruptions, enhanced fitness was detected in the triploid strain. By comparing gene expression across generations, our results indicate that proteostasis restoration is a critical component of rapid adaptation following genome merging in *Chlamydomonas reinhardtii* and possibly other systems.

## INTRODUCTION

Polyploidy, which occurs when cells or organisms possess more than two complete sets of genomes, comes in two types: autopolyploidy, arising from whole genome duplication (WGD), and allopolyploidy, resulting from WGD combined with genome merging, *i.e.* hybridization (Stebbins 1947). It is however often more accurate to describe allo- and autopolyploids as extremes of a continuum along the genetic distance between parental genotypes (Soltis et al. 2010). This ‘mutation’ significantly impacts all biological levels and is prevalent across eukaryotes, but particularly affecting the evolution of angiosperms (Otto and Whitton 2000; Albertin and Marullo 2012; Fox et al. 2020). Indeed, polyploidy is prevalent in contemporary plants, particularly in crops and invasive species, and it seems to confer robustness during environmental stress and upheavals (Fawcett et al. 2009; Soltis et al. 2009; Tate et al. 2009; te Beest et al. 2012; Vanneste et al. 2014; Van de Peer et al. 2017, 2021; Shimizu 2022).

Genome doubling and merging have substantial influences across all levels of cellular biology (Comai 2005; Doyle and Coate 2019; Bomblies 2020). A frequently observed consequence of polyploidy is cell size increase (Otto 2007; Kwak et al. 2017; Doyle and Coate 2019; Bomblies 2020), although this relationship exhibits complex dynamics (Tsukaya 2013). Cell size increase resulting from polyploidization can subsequently affect transcriptome size and transcription (Wu et al. 2010; Marguerat and Bähler 2012; Doyle and Coate 2019). Additionally, genome doubling and, in particular, genome merging have been identified to induce a “genome shock” (McClintock 1984), initiating fast and significant alterations to the genome (Otto 2007). WGD is also expected to alter the balance between the different genomes of plant cells (*i.e.* the stoichiometry between nucleus, plastids and mitochondria, known as cytonuclear stoichiometry), although research on this aspect of polyploidy remains limited (Fernandes Gyorfy et al. 2021).

Genome merging likely has a more profound impact on transcription than genome duplication (Doyle et al. 2008; Chelaifa et al. 2010; Parisod et al. 2010; Spoelhof et al. 2017; Behling et al. 2022). In allopolyploids, gene expression frequently deviates from the additivity hypothesis, which assumes that the gene expression levels are the average of those in the parent species, revealing complex parental legacies (Adams et al. 2003; Osborn et al. 2003; Otto 2003; Adams and Wendel 2005; Auger et al. 2005; Chelaifa et al. 2010; Yoo et al. 2013, 2014). A commonly observed pattern is expression level dominance (ELD), or genome dominance, in which the expression of a given gene is similar to only one of the parents (Rapp et al. 2009; Flagel and Wendel 2010; Grover et al. 2012; Yoo et al. 2013; Combes et al. 2015; Edger et al. 2017; Bird et al. 2018; Li et al. 2018; Wu et al. 2018; Nieto Feliner et al. 2020; Wei et al. 2021). Allopolyploids can also exhibit transgressive expression, where the gene expression is either greater or lesser than that of both progenitors (Rapp et al. 2009; Flagel and Wendel 2010; Yoo et al. 2013). Nonetheless, some allopolyploids exhibit additive expression (Chagué et al. 2010; Chelaifa et al. 2013). Moreover ELD seems influenced by environmental factors (Bardil et al. 2011; Shimizu-Inatsugi et al. 2017), tissue specificity (Li et al. 2014, 2020), and developmental stage (Jia et al. 2022). Gene expression novelty created upon genome merging has been referred to as a ‘transcriptomic shock’ (Buggs et al. 2011). Such alterations might drive the development of novel phenotypes that could be key to the adaptive success of allopolyploids (Hegarty and Hiscock 2009).

Two poorly understood consequences of genome merging on gene expression are genome dosage effects and the evolution of expression. Asymmetric genome inheritance, *via* gene dosage (the number of copies of a gene), could cause asymmetric parental legacies in gene expression in the allopolyploid, as seen *e.g.* in wheat (Qi et al. 2012) and the fish *Cobitis* (Bartoš et al. 2019). It is crucial to track newly formed allopolyploids immediately post-merging to understand the dynamics of gene expression in the first generations after genome merging. Employing transgenerational comparative transcriptomics can elucidate the trends in gene expression following allopolyploidization. Research on wheat presents varied findings regarding post-polyploidization gene expression, as shown in the studies of Chagué *et al*. (2010) and Qi *et al*. (2012). Both studies confirm the immediate alterations in gene expression patterns upon genome merging. However, while Chagué et al. (2010) reported stability within specific gene expression patterns, Qi et al. (2012) observed predominantly stochastic variations. This divergence highlights the need for further investigation to understand the mechanisms behind gene expression patterns in allopolyploids and their evolutionary impacts.

*Chlamydomonas reinhardtii* is a single-celled green alga belonging to the Chlorophyta phylum. This model species is easy to cultivate and manage in the laboratory and has a short generation time. Given these attributes, *C. reinhardtii* serves as a model organism for fundamental research on genetics, photosynthesis, and structural biology (Harris 2001; Sasso et al. 2018; Salomé and Merchant 2019). This microalga species is also valuable in applied research domains, including biopharmaceuticals and biofuel production (Scranton et al. 2015; Kwak et al. 2017). Additionally, its rapid doubling time of eight hours, ease of cultivation, and minimal space requirements make *C. reinhardtii* particularly effective for studying evolution (Bell 1997; Harris 2001; Ratcliff et al. 2013; Bafort et al. 2023).

Here, we employ *C. reinhardtii* to examine the effects of asymmetric genome merging on fitness, expression patterns, and evolutionary changes in gene expression over 425 generations of a laboratory natural selection (LNS) experiment. By crossing a haploid and a diploid strain to create triploid lines, we analyzed the impact using fitness assessment, qPCR, and RNA sequencing at three time-points: the ancestor (G0), generation 225, and generation 425. Our results reveal significant changes in gene expression, with no clear genome dominance, and suggest ‘functional dominance’ from the haploid parent. The study highlights disruptions in cytonuclear balance and potential disturbances in proteostasis, reflecting the complex cellular responses to genome merging. Despite the major nucleo-cytoplasmic changes, the newly-formed triploid strain revealed increased fitness compared to its parental strains.

## MATERIALS AND METHODS

### Strains and experimental conditions

*Chlamydomonas reinhardtii* strains CC-1067 (*arg2*, *mt+*, haploid) and CC-1820 (*arg 7-2*, *mt-*, diploid) were obtained from the Chlamydomonas Resource Center. A triploid strain was formed by fusion of CC-1067 and CC-1820 and complementation of the auxotrophic *arg* mutations, following the method of Ebersold (1967). When not in active cultivation, the strains were preserved on Tris-acetate-phosphate (TAP) agar plates containing arginine under low light conditions (approximately 50 µmol.m^−2^.s^−1^ PPFD) at 22°C. Ploidy levels of the strains were determined and confirmed using polymerase chain reaction (PCR) targeting the mating type locus and flow cytometry, following an adapted version of protocol 6 from Čertnerová and Galbraith (2021).

The triploid strain was bottlenecked to a single cell, followed by its rapid division into five separate, independent lines, initiating the LNS experiment. The experimental lines were cultivated in 5 mL arginine-supplemented TAP medium within Erlenmeyer flasks at 23°C, under a light intensity of 150 µmol.m ^−2^.s^−1^ PPFD and agitated at a rate of 150 rpm. Routine transfers of the lines were performed, consisting of transferring 5% of the volume into fresh medium, occurring once or twice per week.

Evolving lines were cryopreserved at regular time intervals to allow future comparative analyses with evolving lines. After 99 transfers, we thawed the ancestral strains (CC-1067, CC-1820, and the triploid line following bottlenecking and prior to line division, thereafter named 1N parent, 2N parent, and 3N G0, respectively) and the experimental triploid lines from the 52^nd^ transfer (3N G225).

The number of generations was calculated using the formula:

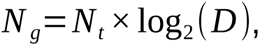

where *N _g_* represents the number of mitotic generations, *N_t_* denotes the number of transfers, and *D* is the dilution factor used at each transfer. For this experiment, the dilution factor *D* was set at 20. Consequently, lines cryopreserved after undergoing 52 transfers, resulted in a calculated generation number of 52 *×*log_2_(20) *≈* 225. Similarly, lines harvested after 99 transfers, had a generation number calculated as 99 *×* log_2_(20)*≈* 425.

Thawed and evolving LNS lines were subjected to periodic conditions (12 hours at 28°C, 220 µmol.m^−2^.s^−1^ PPFD: 12 hours at 18°C, dark) to synchronize the cultures prior to cell harvest, flow cytometry and growth assays. The conditions for synchronization were ascertained by following the methodologies stipulated by Hlavová et al. (2016) and Angstenberger et al. (2020), supplemented with a process of laboratory trial and error. See Table 1 for a summary of all experimental strains. Once sufficient synchronization in the cell cycle was observed for all lines, as confirmed through microscopy, each experimental line was divided into six replicates, randomized, and cultivated for three days under the same conditions as in the LNS experiment and in the same incubator. Seventy-two hours post-inoculation, 2 mL aliquots of each independent line were centrifuged to remove liquid culture, and the resulting samples were flash-frozen in liquid nitrogen. These samples were subsequently stored at -80°C until RNA extraction. Concurrently, an additional 1 mL sample was collected from each line for cell count (Multisizer 3 Coulter Counter).

### Fitness assays

Every experimental line was diluted to 1:25, taking it below the detection threshold of our plate reader, and divided into 12 replicates, which were randomly distributed onto 96-well plates (ref. 655098, Greiner Bio-One). The optical density was measured approximately every 12 hours over a span of eight days using a VICTOR™ X Multilabel Plate Reader (PerkinElmer, Inc.). We applied the Baranyi-Roberts equation to model growth curves, facilitating estimates of the maximum growth rate and population size (Baranyi and Roberts 1994).

We applied the following strategy to detect significant differences between the experimental lines for MGR. First, to ascertain the homoscedasticity and normality of the residual distributions associated with MGR, Levene’s test and the Shapiro-Wilk test were employed respectively, as implemented in R. If the normality assumption was satisfied, t-tests were used to detect difference. If the normality assumptions were not met, Mann-Whitney U tests were employed. To account for multiple comparisons, p-values were adjusted using the Benjamini-Hochberg correction to control the false discovery rate.

### RNA extraction and RNA-Seq

Total RNA was extracted using a Qiagen RNeasy® Plant Mini Kit and treated with RNase-free DNase (Qiagen, Stanford, CA, USA). The assessment of total RNA quality and quantity was conducted utilizing a NanoDrop™ 2000c spectrophotometer (Thermo Fisher Scientific) and Bioanalyzer RNA6000 (Agilent Technologies). Library preparation and sequencing were conducted by the sequencing platform *NXTGNT* (http://www.nxtgnt.com/). cDNA libraries were generated using a Quantseq™ 3′ mRNA-Seq library preparation kit (Lexogen) in accordance with the manufacturer’s guidelines. The Illumina NextSeq 500 was utilized for sequencing, yielding an average of 11.3M 76 bp single-end reads per sample (range: 8.7M-20.1M reads). Following Quality Control (FastqC), Unique Molecular Identifiers (UMIs) were extracted using ’umi_tools extract’ (Smith et al. 2017) and the reads were trimmed utilizing ’bbduk’ (Bushnell 2014). Reads from identical libraries (processed in distinct runs and lanes) were amalgamated prior to ’hisat2’ mapping (Kim et al. 2019) onto the *C. reinhardtii* genome version 6 (Craig et al. 2023). On average, 9M reads mapped one time to the reference genome (range: 6.9M-16.6M). PCR duplicates were eliminated by employing the UMIs and ’umi_tools dedup’ (Smith et al. 2017). Mapped reads were quantified using the Python package HTSeq (Anders et al. 2015).

### Differential gene expression analysis

Differential expression (DE) analysis was performed with the Bioconductor package *HybridExpress* on count data normalized by library size (Almeida-Silva et al. 2024). This package facilitated the calculation of midparent values (MPV) for gene expression, exploratory data analysis, DEG analysis, gene categorization into different expression patterns (see paragraph below), and KEGG pathway overrepresentation analysis. Differential expression was determined using an adjusted P-value (Benjamini-Hochberg correction) threshold of *P*<0.01. Differentially expressed genes were classified into categories and classes of expression patterns as developed by Rapp et al. (2009) and as implemented in the R package *HybridExpress*.

### Inference and analysis of gene coexpression networks

Gene coexpression networks (GCNs) were inferred and analyzed with the Bioconductor package BioNERO (Almeida-Silva and Venancio 2022). Prior to network inference, count data were filtered to remove genes with less than one count in at least 50% of the samples, followed by variance-stabilizing transformation (Love et al. 2014). We inferred a single GCN containing all samples, and five strain-specific GCNs containing only samples from the triploid ancestral line and evolved lines at generations 225 and 425. GCNs were inferred using biweight midcorrelations and the signed hybrid approach. Module preservation statistics were calculated using the NetRep method (Ritchie et al. 2016) implemented in BioNERO, with 1000 permutations. Modules in the reference network (Line 1) were considered as preserved in other networks if at least five (of seven) preservation statistics were significant (*P*<0.05).

### Quantification of nuclear and cytosolic DNA by qPCR

The analysis of relative DNA content was conducted using quantitative PCR (qPCR). Cells were collected from 0.5 mL cultures through centrifugation at 1000 g for 5 minutes, immediately frozen in liquid nitrogen, and then stored at -80°C for DNA extraction. To mitigate the impact of cell cycle variations, which can affect genome balance in *C. reinhardtii* (Kabeya and Miyagishima 2013), samples from each strain were collected at six different times across two days (10 AM, 2 PM, and 5 PM). DNA extraction was performed using the Omniprep^TM^ for Plant DNA extraction kit (G-Biosciences), and the extracted DNA was diluted to a final concentration of 0.1 ng.μL^-1^. The LightCycler 480 SYBR Green I Master (Roche Diagnostics) was used for qPCR, and the reactions were run on a LightCycler® 480 Instrument II (Roche Diagnostics). Primers targeting the nuclear DNA (EIF1A gene), chloroplast DNA (rbcL gene), and mitochondrial DNA (nd1 gene) were employed, with sequences 5’-CATTGTGGAGCCGCCATTTC-3’ and 5’-GGCTGCTTGCATTTGCTTCC-3’ for EIF1A, 5’-GTCGTGACCTTGCTCGTGAA-3’ and 5’-TCTGGAGACCATTTACAAGCTGAA-3’ for rbcL, and 5’-GATCTACCAGAGGCTGAGTTG-3’ and 5’-TTTAAGGGCGCTGAAGCCAC-3’ for nd1. Each sample underwent qPCR analysis in triplicate to ensure accuracy, with the resulting data points averaged for subsequent analysis. Prior to relative quantification, primer efficiency was validated through a dilution series. Relative DNA content quantification between samples was performed utilizing the double delta Ct method, employing the haploid strain as the baseline for comparison.

## RESULTS

### Genome merging slightly increases fitness and triploid lines show potential adaptation to experimental conditions

We assessed the fitness of experimental lines using a growth assay to measure Maximum Growth Rate (MGR) as a fitness proxy (Fig. 1). Detailed growth curves can be visualized in Supplementary Figure S1. The newly-formed triploid (3N G0) showed significantly higher MGR compare to its diploid parent and haploid parent ( *P*<0.001 and *P*=0.035, respectively, t-tests adjusted using Benjamini-Hochberg correction). We also compared the MGR of evolved triploid lines at generation 225 (G225) with both the ancestral triploid (3N G0) and with these lines at G425 (Fig. 1).

**Figure 1.**
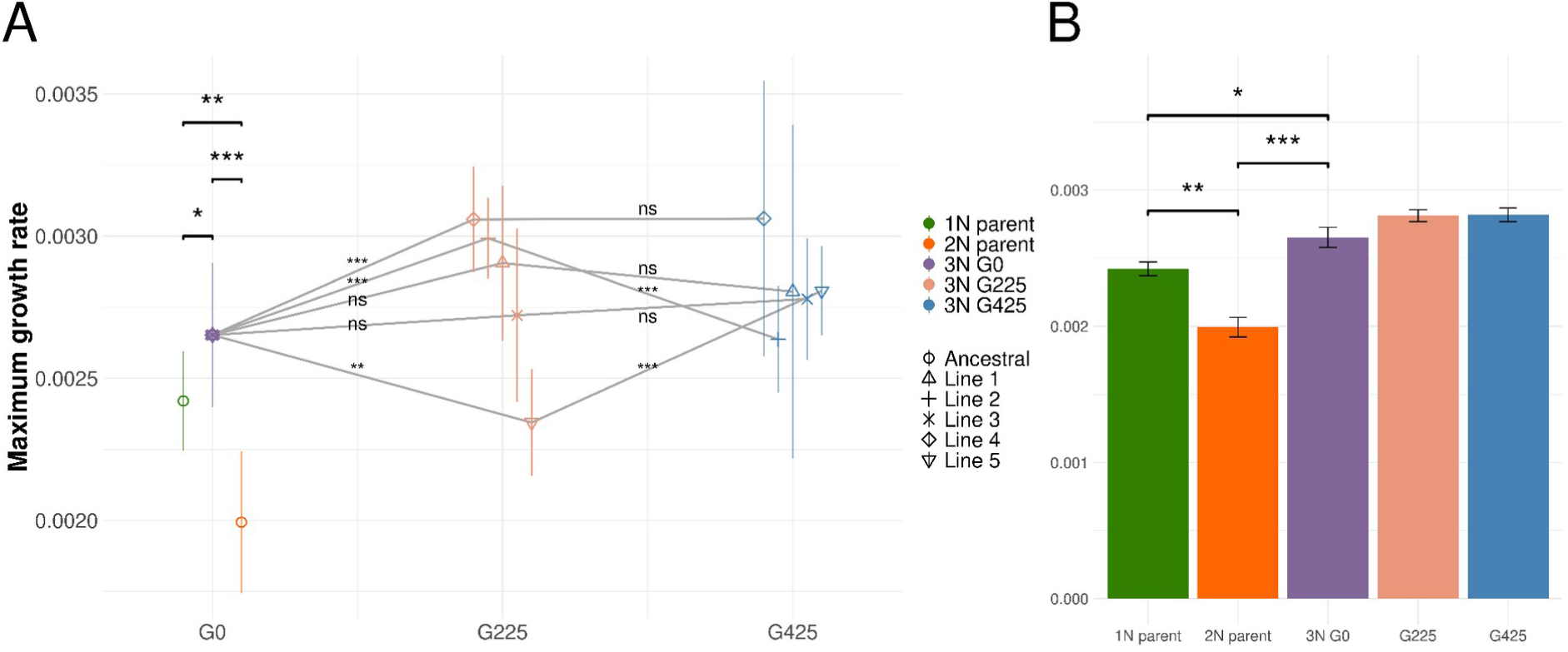
Fitness assessment of experimental lines using maximum growth rate (MGR). **A**: Mean MGR of the experimental lines across three generations (G0, G225 and G425). Error bars represent standard deviation. Statistical significance of differences between lines, determined by t-tests or Mann-Whitney U tests based on data distribution, is indicated above comparisons (‘ns’ for not significant, ‘*’ for p<0.05, ‘**’ for p<0.01, ‘***’ for p<0.001). **B**: Mean MGR of parental strains, ancestral triploid and triploid lines pooled at generation time-points (G225 and G425). Error bars represent standard errors. Presented strains include haploid CC-1067 (1N parent, green), diploid CC-1820 (2N parent, orange), ancestral triploid (3N G0, purple), and a pooled representation of evolved triploid lines (Lines 1-5) at generations 225 (salmon pink) and 425 (blue).

MGR of the five independent triploid lines were pooled and averaged by generations for comparison. We observed a slight increase in MGR from G0 to G225, followed by stabilization from G225 to G425.

### Persistent and evolving gene expression in triploids across generations

Significant differential expression (DE) was detected between the two parental strains, with 6,096 genes—approximately 36.1% of the total gene count—being differentially expressed, using a p-value threshold of 0.01 (Fig. 2). Substantial DE levels were also evident between the parental strains and the initial triploid (3N G0), showcasing similar differentially expressed gene (DEG) proportions for the haploid and diploid parent, at 26.8% and 29.7%, respectively (Fig. 2A). 21.4% of genes demonstrated DE between the initial triploid and the MPV (Fig. 2A). A principal component analysis (PCA) of gene expression levels (Fig. 3) clearly separates the triploid derivative from the MPV and each parental strain, indicating distinct expression profiles. The analysis reveals that the triploid line does not manifest intermediate expression levels typical of parental additivity. Instead, it forms a well-defined cluster, distinct from both the parental strains and the MPV. Furthermore, while replicates within the same lineage number and LNS lines cluster tightly, G225 and G425 show no significant separation. However, LNS lines from G225 subtly tend to cluster closer to the ancestral state compared to those from G425.

**Figure 2.**
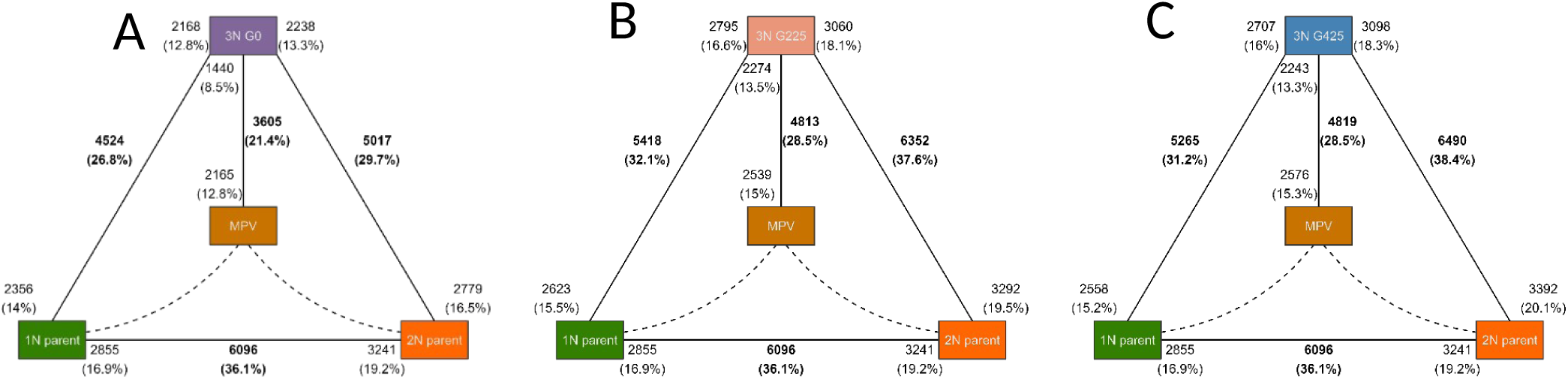
Differential gene expression among ancestral parental strains and triploid lines. The haploid parent CC-1067 (“1N parent”) is shown in green, the diploid parent CC-1820 (“2N parent”) in orange, and the *in silico* midparent (“MPV”), representing the averaged expression profile of the two parents, in brown-orange. Each panel shows the total count and proportion of differentially expressed genes, indicating the direction of up-regulation for each comparison. **(A)** Comparisons with triploid progeny at generation 0 (3N G0). **(B)** Comparisons with triploid lines at generation 225 (3N G225). **(C)** Comparisons with triploid lines at generation 425 (3N G425).

**Figure 3.**
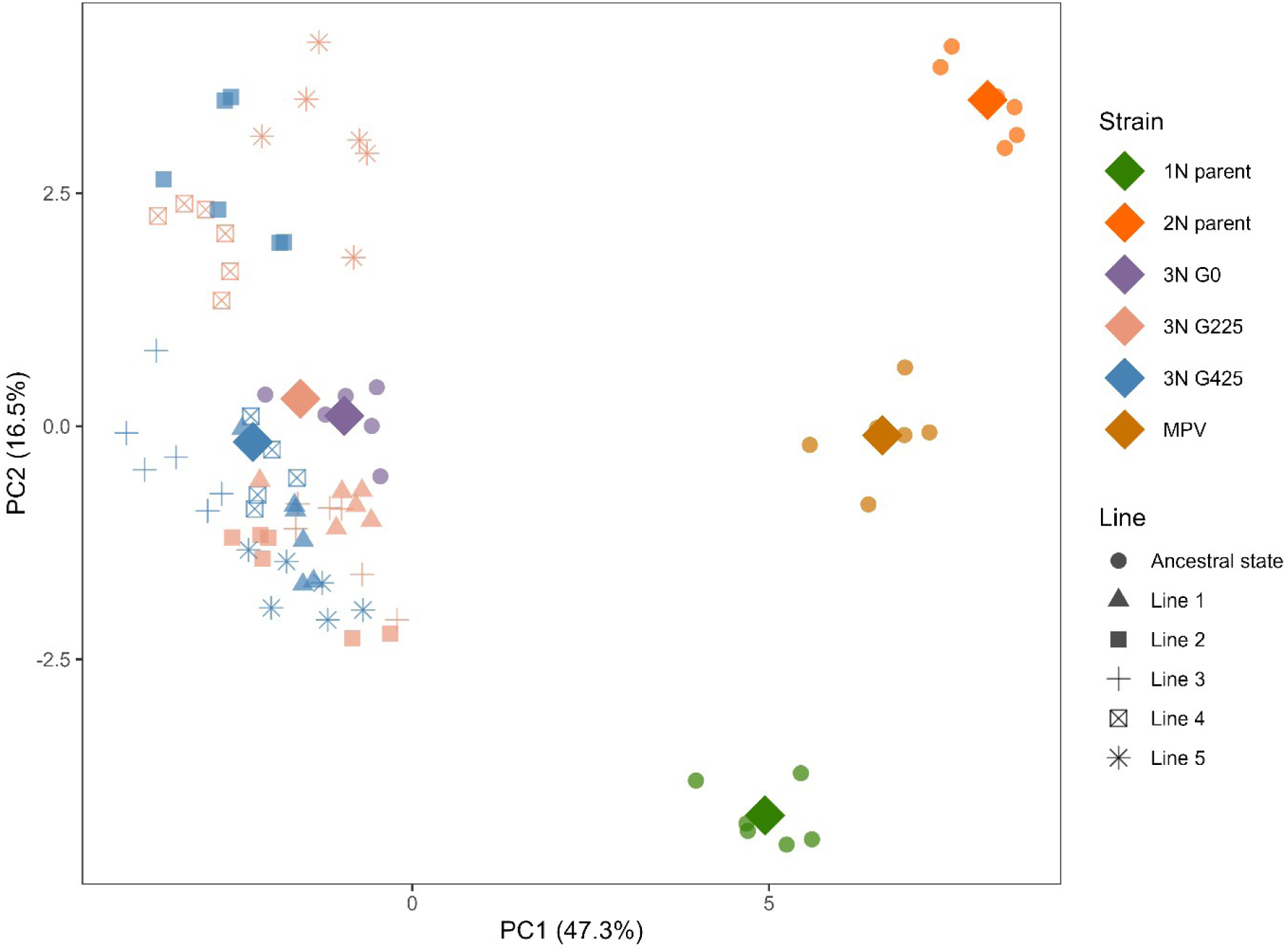
Principal component analysis of differential expression levels. Each individual replicate is depicted by a small, slightly transparent dot, with the shape varying according to the associated line. Group aggregates are denoted by larger, opaque rhombus symbols. The ancestral strains, comprising the haploid parent, the diploid parent, the triploid derivative, and the midparent values (MPV), are distinguished by color codes: green, orange, purple, and brown-orange, respectively. Evolved triploids are illustrated using distinct shapes, contingent on the lines, and unique colors, contingent on the generation: generation 225 is represented in salmon pink, while generation 425 is delineated in blue.

Genes exhibiting DE between at least one pair within the trio —Haploid parent, Diploid parent, and Triploid derivative—were classified into 12 expression patterns as delineated by Rapp et al. (2009) using the *HybridExpress* function *expression_partitioning* (Almeida-Silva et al. 2024). These patterns were further grouped into five broad classes: transgressive up-regulation (UP), transgressive down-regulation (DOWN), additivity (ADDITIVE), expression level dominance (ELD) toward the haploid parent (ELD1), and ELD toward the diploid parent (ELD2). Gene expression pattern classification was applied to the ancestral triploid (G0), as well as collectively across the five independently evolving triploid lines at G225 and G425. By pooling the data from these lines, we aimed to identify expression patterns indicative of selective pressures, thereby minimizing the potential confounding effects of genetic drift. Consistent with preceding DE results, only 9.73% (603) of the DEGs showed additive expression at G0, exhibiting levels intermediate between the parental strains. This number rose to ∼15.5% at both G225 and G425 (1094 and 1093, respectively). Transgressive expression constituted a substantial fraction of DEGs, comprising 9.23% (572) of up-regulated genes and 14.09% (873) of downregulated genes at G0. Upregulated gene numbers increased to ∼12% at both G225 and G425 (852 and 854, respectively). The downregulated gene fraction slightly decreased with 13.47% (953) and 13.88% (980) genes at G225 and G425, respectively. Remarkably, at G0, around two-thirds (66.96%, 4150 genes) of DEGs showed ELD. At G0, genes showing ELD1 and ELD2 were at proportions of 36.22% (2245) and 30.74% (1905) respectively. Although the proportion of ELD1 genes remained stable, the absolute number of these genes increased from 2245 at G0, to 2416 at G225, and finally to 2478 at G425. ELD2 genes declined to 24.86% (1758) and 23.45% (1656) at G225 and G425, respectively.

A notably high proportion of genes share constant expression patterns across three generations, particularly for ELD, ranging from 26% for genes consistently up-regulated to 45% for genes exhibiting ELD1. These genes, maintaining consistent class categorization across generations, are henceforth termed ’persistent genes’ (Fig. 4B).

**Figure 4.**
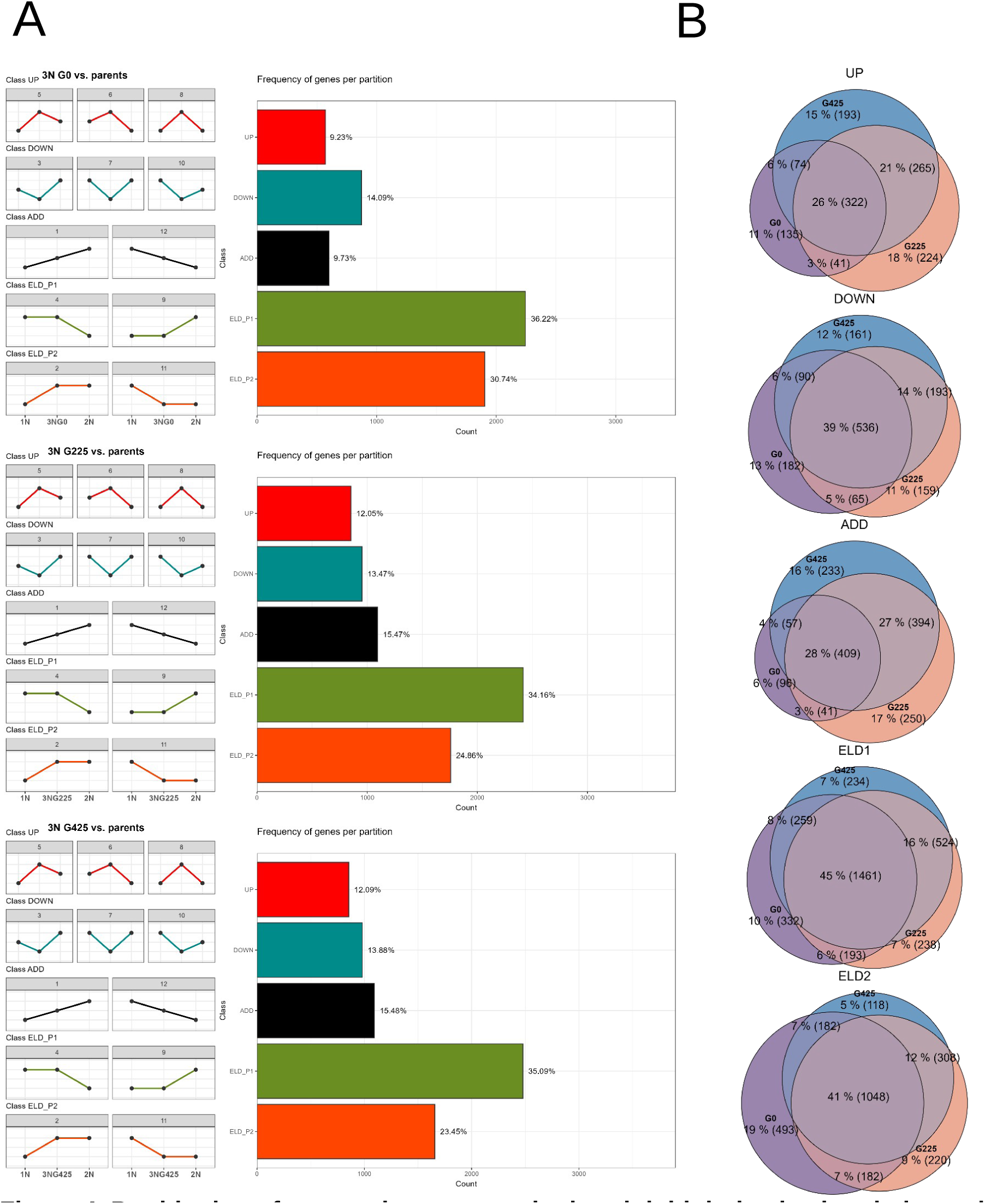
Partitioning of expression patterns in the triploid derivative in relation to its haploid and diploid progenitors at generation 0 (G0), generation 225 (G225), and generation 425 (G425). The differentially expressed genes are binned in five distinct expression patterns: transgressive upregulation (UP, in red), transgressive downregulation (DOWN, in blue), additivity (ADDITIVE, in black), expression level dominance toward the haploid parent (ELD_P1 or ELD1, in green), and expression level dominance toward the diploid parent (ELD_P2 or ELD2, in orange). **(A)** Left: graphical representations detailing the 12 potential expression patterns observed between the two parental strains and their derivative, binned in five groups; Right: bar plots showing the number and fraction of differentially expressed genes that fall in the five possible expression patterns in the triploid at G0, G225, and G425 (from top to bottom, respectively). **(B)** Euler diagrams illustrating the number and percentage of genes consistently represented within the five expression patterns across G0, G225, and G425 in the triploid progeny lines.

To follow the evolution of expression level after genome merging, genes that showed DE between at least one pair within the trio—Ancestral triploid (3N G0), triploid LNS lines at G225 (3N G225), and triploid LNS lines at G425 (3N G425)— were sorted into the analogous 12 expression patterns employed for the three ancestral strains in the preceding analysis (Fig. 5). This approach enabled the identification of any significant changes in expression levels throughout the duration of the LNS experiment. A total of 1040 genes demonstrated DE between at least one of the comparisons. A substantial 81.7% of the DEGs fell into categories 2, 11, 7, and 8, leaving the remaining eight categories with considerably fewer genes. Categories 2 and 11 (286 and 159 genes, respectively) corresponded to genes that underwent significant changes in expression levels within the initial 225 generations, subsequently stabilizing. Categories 7 and 8 (175 and 230 genes, respectively) corresponded to genes that manifested similar expression levels between G0 and G425, but significantly different expression at G225.

**Figure 5.**
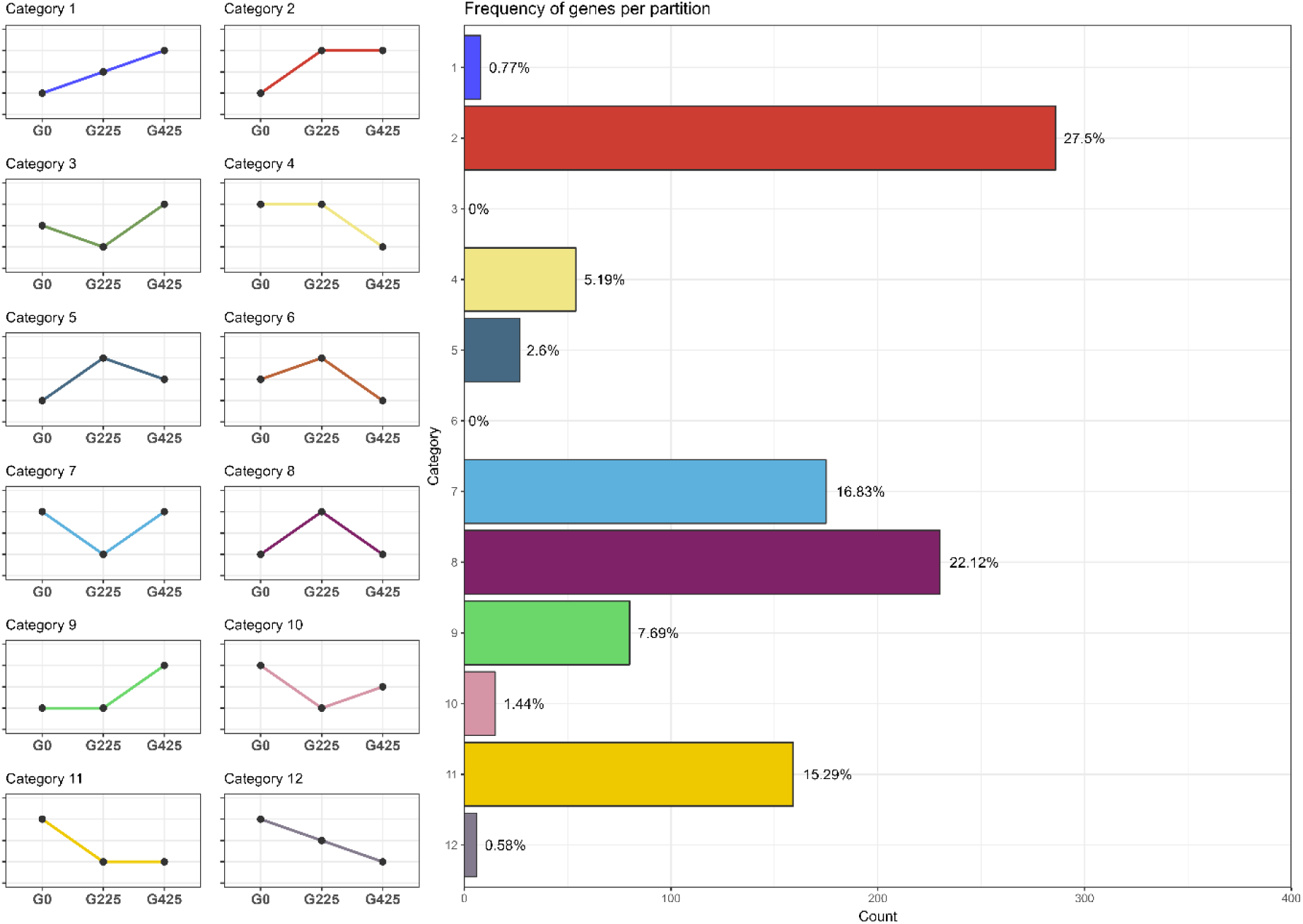
Partitioning of patterns of expression level evolution in the triploid lines. Left: graphical representations detailing the 12 possible expression categories observed between the three generations of the triploid lines; Right: bar plots showing the number and fraction of differentially expressed genes that fall in the 12 possible categories (right).

### Enrichment analysis reveals immediate and evolutionary consequences to genome merging in gene expression

We examined enrichment of KEGG metabolic pathways in genes classified in the five expression patterns observed consistently at G0, G225, and G425, termed ‘persistent genes’ (Fig. 4B). Upregulated genes (UP) demonstrated no significant enrichment in any metabolic pathways. Conversely, downregulated genes (DOWN) exhibited substantial enrichment, particularly in KEGG pathways associated with the chloroplast and mitochondria, including “Photosynthesis” and the “Krebs cycle” (Supplementary Table 1). Additive genes (ADD) showed no pathway enrichment overall. ELD1 genes were primarily enriched in pathways related to ribosomes and the metabolism of proteins and amino acids, as outlined in Supplementary Table 2. Similar to the UP and ADD classes, genes with 2N Expression Level Dominance (ELD2) displayed no significant pathway enrichment.

We also analyzed the overrepresentation of gene ontology (GO) terms in genes within the different categories of evolution (Fig. 5). Among the 12 categories, only categories 2 and 11 showed significant enrichment. Category 2, which includes genes that showed a rapid increase in expression levels during the experiment, displayed enrichment in terms associated with “autophagy” and “protein catabolic process” (Supplementary Table 3). Category 11, representing genes that experienced a rapid decrease in expression, showed enrichment in “translation” and “peptide biosynthetic process” (Supplementary Table 4). This suggested an evolution of the peptide anabolism and catabolism processes in the triploid lines, potentially caused by an initial disruption after genome merging. To investigate this, we examined KEGG pathway enrichment in DOWN and UP genes in the ancestral triploid at G0. Notably, UP genes showed significant enrichment in “Ribosome biogenesis in eukaryotes,” while DOWN genes showed significant enrichment in “Proteasome.” indicative of a disruption of proteostasis following genome merging.

### qPCR confirms disruption of cytonuclear stoichiometry in triploids

We employed qPCR to quantify the relative DNA content from nuclear, plastid, and mitochondrial genomes in the triploid (3N G0) and haploid *mt+* parent strains (1N parent). This approach was motivated by our preliminary findings, which indicated a downregulation of genes associated with mitochondrial and chloroplast functions in the triploid strain. Given Chlamydomonas’s uniparental inheritance of cytosolic genomes, we posited that the balance between cytosolic (chloroplast and mitochondria) and nuclear genomes might be altered in the triploid compared to its haploid counterpart. qPCR results confirmed this hypothesis. The ratio of chloroplast DNA (cpDNA) to nuclear DNA (nuDNA) and mitochondrial DNA (mtDNA) to nuDNA in the triploid strain was significantly reduced, approximately threefold lower, compared to the haploid parent. Specifically, the cpDNA/nuDNA and mtDNA/nuDNA ratios in the triploid were 0.345 and 0.325, respectively. These findings suggest a substantial shift in genome balance, with a marked decrease in the relative content of cpDNA and mtDNA in the triploid strain.

### Gene coexpression networks reveal temporal changes in biological processes

We inferred a gene coexpression network (GCN) with all samples using BioNERO (Almeida-Silva and Venancio 2022) and identified 62 modules, of which 15 were enriched in genes associated with Gene Ontology terms and/or KEGG pathways (Fig. 6A; Sup. Fig. S2). As per BioNERO’s default behavior, coexpression modules are represented by different color names. Module *blue* contained genes involved in the biosynthesis of secondary metabolites and carbon metabolism, with decreased expression levels in triploid G0 (Fig. 6B). Genes in modules *blue2* and *darkseagreen3* were associated with response to osmotic stress, and non-coding RNAs (ncRNA processing, gene silencing by miRNAs, and histone methylation), respectively, and their expression in triploid G0 corresponded to the mean of the 1N and 2N parents (Fig. 6B). Genes in module *darkgreen* were involved in cell cycle, and displayed dramatically lower expression levels in the 1N parent and triploid G0. Module *darkslateblue* contained genes involved in rRNA maturation and regulation of ribosome biogenesis, and they displayed increased expression levels in triploid G0, with ever-increasing expression levels over time in evolved lines.

**Figure 6.**
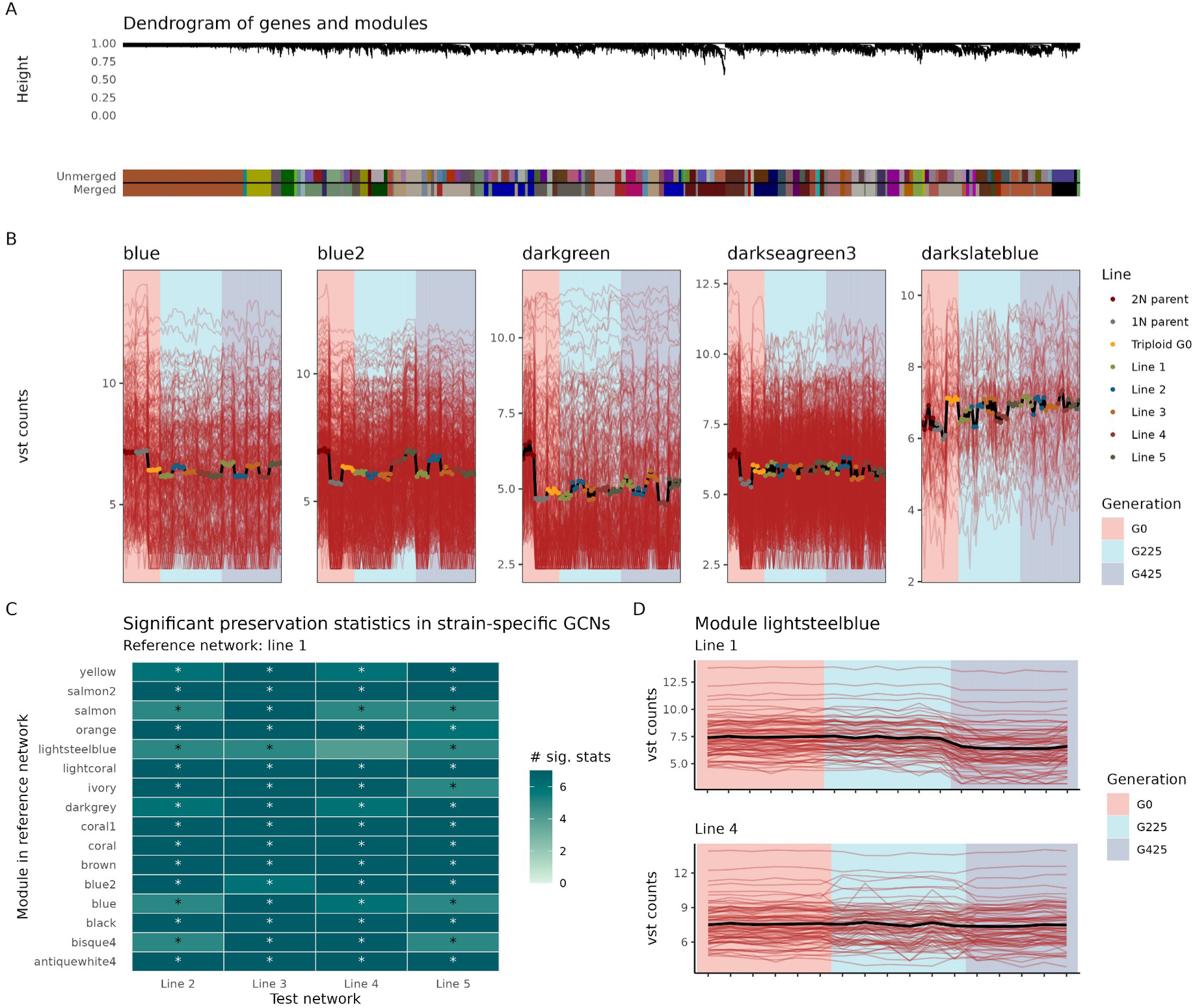
Gene coexpression network analyses. A. Dendrogram of genes and modules obtained with BioNERO. Modules with correlations between eigengenes >0.8 were merged into a single larger module to remove redundancy. **B.** Expression profiles of selected modules enriched in functional terms (Gene Ontology and/or KEGG pathways). Expression levels are represented as variance-stabilized count data (*i.e.*, vst counts). **C.** Significant network preservation statistics between reference and test strain-specific coexpression networks (GCNs). Statistics were obtained by comparing modules in the reference strain-specific gene coexpression network (line 1) with all other strain-specific networks. All preservation statistics in the NetRep algorithm were used. Asterisks indicate modules that had at least five significant preservation statistics. **D.** Expression profiles of the genes in module *lightsteelblue* (reference network) in lines 1 and 4. The module *lightsteelblue* was the only module in the reference network that was not preserved in other test networks. The line plots indicate that expression divergence between Line 1 and Line 4 occurs after generation 425.

Further, we hypothesized whether there is an association between a gene’s expression-based class (*i.e.*, UP, DOWN, ADD, ELD1, and ELD2; see previous sections) and its degree in the GCN (*i.e.*, sum of all edge weights). We observed that genes in classes UP, DOWN, and ADD were overrepresented in hubs ( *P*<0.001). Based on numerous reports on the association between hub genes and essentiality in a cell, with hub gene knockouts leading embryo lethality (Jeong et al. 2001; Yu et al. 2004; Zotenko et al. 2008; Song et al. 2015; Almeida-Silva et al. 2020), this finding suggests that genes in these classes have a more prominent role in the organism’s fitness.

### Most genes displayed preserved expression levels across different evolved lines

We observed some variation in expression levels across different lines within the same generation (Fig. 6B). To test whether different lines had divergent expression profiles over generations, we inferred GCNs separately for each line (hereafter referred to as ‘strain-specific GCNs’ or ‘ssGCNs’). We then calculated module preservation statistics between a reference ssGCN (for Line 1) and all other test ssGCNs (Lines 2, 3, 4, and 5) using preservation statistics implemented in the NetRep algorithm (see Materials and Methods for details). We observed that all modules in the reference ssGCN were preserved in the ssGCNs for nearly all test ssGCNs, except for module *lightsteelblue* in the ssGCN for Line 4 (Fig. 6C). After further investigation, we found that divergence in expression profiles between Lines 1 and 4 occurred after generation 425 (Fig. 6D), with a decrease in expression levels in Line 1, but not in Line 4. Functional enrichment analyses revealed no enriched terms for genes in module *lightsteelblue*.

## DISCUSSION

### Asymmetric genome merging causes transcriptomic shock

Genome merger can produce gene expression patterns that show intermediate levels of the parent species, as suggested by the additivity hypothesis (Buggs et al. 2014; Yoo et al. 2014). Indeed, many homoploid hybrids and allopolyploids exhibit predominantly additive gene expression relative to their parents (Chelaifa et al. 2013; Zhang et al. 2016; Bartoš et al. 2019; Zhou et al. 2019). However, examples of “transcriptomic shock”, characterized by extensive non-additive gene expression, are also well documented (Hegarty et al. 2006; Wang et al. 2006; Flagel and Wendel 2010; Wu et al. 2018; Li et al. 2020). Such shocks lead to novel expression patterns, introducing phenotypic variations that could drive adaptation (Mable 2013; Van de Peer et al. 2021).

Our RNA-Seq experiment uncovered a pronounced transcriptomic shock in our newly formed triploid *C. reinhardtii* line, characterized by large differences in gene expression and unique gene expression patterns. The extent of this shock is surprising, as it typically occurs when genomes from different species are merged, whereas in this case, the genomes of two strains from the same species were combined. However, the parental strains, despite belonging to the same species, exhibited substantial differences in gene expression (Fig. 2), which might explain the unexpected shock observed in the triploid (Zhang et al. 2019). This marked discrepancy in expression might be attributed to the differing ploidy levels (haploid *vs*. diploid) and haplotype differences. We note that mutations could have disrupted the expression patterns of the diploid, as this strain was exposed to a mutagenic agent that potentially caused its diploidization (Loppes 1969). The transcriptomic shock in the triploid appears to have occurred immediately following genome merging, as evidenced by its presence in the ancestral triploid strain (G0), and it has persisted over 425 subsequent generations. Although variation exists, both the patterns of gene expression and the specific genes involved showed a tendency for inheritance (Fig. 4), contrasting with previous results in newly-formed allohexaploid wheat (Qi et al. 2012). These results provide a compelling example of novel and heritable expression patterns emerging very rapidly after genome merging within a single sexual generation. Interestingly, the triploid lines showed an increase in fitness, approximated by MGR, compared to both parental strains (Fig. 1). This suggests that the observed transcriptomic shock, rather than detrimentally affecting fitness, potentially contributed to increased it under our laboratory conditions.

### Complex parental legacy observed in the triploid lines

Strong gene-level dominance was evident, as approximately two-thirds of the DE genes exhibited ELD towards either the haploid or the diploid parent. Notably, despite asymmetric genome inheritance, the triploid progeny did not exhibit genome-wide dominance favoring the diploid parent (*i.e.* ELD2). Moreover, the number of genes demonstrating ELD1 slightly outnumbered those showing ELD2 at G0. This bias towards the haploid parent appeared to intensify in subsequent generations (Fig. 4A), suggesting a potential ongoing parental dominance from the haploid strain.

Additionally, the overrepresentation analyses of the ELD1 and ELD2 gene sets revealed a distinct contrast in biological roles. Specifically, ELD1 genes were enriched in five KEGG pathways, predominantly those involved in amino acid metabolism, whereas ELD2 genes did not show enrichment in any pathway. Considering these observations—the increasing proportion of ELD1 genes relative to ELD2 genes, the similar fitness levels of the triploid and the haploid parent, and the results of the enrichment analysis—we conclude that the triploid potentially exhibits functional dominance towards its haploid parent, despite the unfavorable imbalanced genome inheritance. It is conceivable that this haploid dominance could be due to the higher fitness of the haploid parent relative to the diploid parents in our laboratory conditions (Fig. 1). Indeed, during the LNS experiment, these fitness advantages could have led to a selective pressure favoring traits associated with the haploid genome. Consequently, the prevalence of haploid dominance in the triploid could be an adaptive response, optimizing the triploid’s metabolism to enhance growth under our experimental conditions.

These results align with numerous prior studies showing that allopolyploids often exhibit dominance at the gene expression levels towards one of the parental species (Rapp et al. 2009; Li et al. 2020; Glombik et al. 2021). This phenomenon has been extensively reviewed in the literature, highlighting its prevalence and significance in allopolyploid evolution (De Smet and Van de Peer 2012; Grover et al. 2012; Buggs et al. 2014; Yoo et al. 2014; Wendel et al. 2018). However, the observation that the dominant genome is haploid rather than diploid presents a surprising deviation from previous studies on resynthesized allohexaploid wheat, which predominantly demonstrated a dosage effect influencing expression level dominance (Qi et al. 2012; Li et al. 2014). This deviation could be influenced by external conditions, which significantly affect the parental legacy of gene expression in allopolyploids (Bardil et al. 2011; Shimizu-Inatsugi et al. 2017). Although our strains were cultivated under optimal conditions, the observed “haploid functional dominance” may be due to the superior fitness of the haploid parent in these specific conditions. Further experimentation including genome and epigenome sequencing is needed to confirm these findings and to explore the underlying mechanisms of this dominance.

### Asymmetric genome merging leads to major disruptions of cytonuclear stoichiometry and proteostasis

Enrichment of KEGG pathways and GO term gave insights into the consequences of genome merging for the cell biology of the new triploid strain. The significant presence of KEGG pathways linked to photosynthesis and carbon metabolism in downregulated genes suggested a disruption of the cytonuclear stoichiometry in the triploid (Sup. Table 1). Additionally, our coexpression analysis shows that the module *blue* containing genes involved in carbon metabolism displayed decreased expression level in triploid G0. Given the uniparental inheritance of organellar (chloroplast and mitochondria) genomes in *C. reinhardtii* (Grant et al. 1980; Nakamura 2010), we hypothesized that these enrichment outcomes were caused by a change of the relative copy number of nuclear, mitochondrial and plastid genomes in the triploid. We confirmed this result by qPCR, showing that the ratios cpDNA/nuDNA and mtDNA/nuDNA were approximately ⅓ of those of the haploid parent. This finding is surprising, as *C. reinhardtii* typically increases its chloroplast DNA content with ploidy level (Whiteway and Lee 1977), a trend also observed in *Arabidopsis* autopolyploids (Fernandes Gyorfy et al. 2021). Although normal mating processes predominantly result in maternal inheritance (*mt+*) of chloroplasts (Burton et al. 1979; Kuroiwa et al. 1982) and paternal inheritance (*mt-*) of mitochondria (Nakamura 2010), the inheritance patterns of the organellar genomes in our triploid remain unclear. This is particularly relevant for our triploid strains, as they have not undergone zygospore formation, making the inheritance patterns of chloroplasts even more unpredictable (Gillham 1969). Future genomic sequencing and analysis will be crucial to elucidate these patterns.

Growth assay results (Fig. 1) indicate increased MGR despite the cytonuclear disruption under optimal growing conditions. Similarly, GO enrichment of genes with significant expression changes after genome merging do not suggest any adaptation to this new cytonuclear stoichiometry, such as increased expression of organellar genes (Sup. Tables 3 and 4). This resilience aligns with recent findings on the robustness of cytonuclear interactions following disruptions in allopolyploid angiosperms (Sloan et al. 2024). The minimal impact on fitness could also be attributed to the use of TAP medium, which contains acetate—a carbon source that *C. reinhardtii* can metabolize heterotrophically—possibly mitigating the effects of this disruption on growth (Heifetz et al. 2000).

Evidence on the regulation of organellar pathways in polyploids is limited. Li et al. (2020) observed a downregulation in photosynthesis-related pathways in natural allotetraploid *Brassica napus*, aligning with our findings. Contrarily, numerous studies report an increase in photosynthetic rates, chloroplast density, and chlorophyll content in both established and newly synthesized allopolyploids (Warner and Edwards 1993; Vyas et al. 2007; Coate et al. 2012; Ilut et al. 2012), yet these studies rarely explore the corresponding gene expression levels. However, Coate and Doyle (2013) noted increased expression of certain photosynthesis-related genes, while Forsythe et al. (2022) found that established polyploid plants preserved cytonuclear expression ratios, demonstrating their capacity to adapt to cytonuclear disruptions. Similarly, newly formed autotetraploids of *Festuca pratensis* and *Lolium multiflorum*, induced by colchicine, increased their chloroplast and chloroplast genome copy numbers by approximately twofold to compensate for disrupted cytonuclear stoichiometry, with no significant differences in nuclear or chloroplast gene expression levels (Shahbazi et al. 2024). These contrasting observations underscore the complex effects that genome merging and doubling may have on cytonuclear stoichiometry and/or the regulation of photosynthesis genes (Grover et al. 2022).

The genome merging in the triploid strain notably led to a downregulation of genes involved in protein degradation and an upregulation of those linked to protein biosynthesis, indicating an initial disruption of protein homeostasis (proteostasis). Gene expression analysis post-merging revealed a distinct pattern: significant early changes (between G0 and G225) that later stabilized (between G225 and G425; Fig. 5). GO enrichment analysis indicated that genes which rapidly decreased in expression were primarily associated with protein biosynthesis (Supplementary Table 4), whereas genes with increased expression were linked to protein catabolism (Supplementary Table 3). These findings suggest that the evolution of gene expression in the triploid lines was predominantly driven by selection pressures aimed at restoring proteostasis. As shown in yeast (Lu et al. 2016), excessive protein production appears to be a major intrinsic stress of neopolyploidization, suggesting that restoring proteostasis is a crucial adaptation to polyploidy. However, our understanding of the impact of polyploidy on the proteome remains limited, necessitating further research to fully explore its effects (Soltis et al. 2016; Doyle and Coate 2019).

### Concluding remarks

Our study leverages *C. reinhardtii*, a unicellular green alga closely related to angiosperms, as a unique model to explore the cellular and evolutionary implications of polyploidy (Bafort et al. 2023). This system allows for detailed examination of both the immediate cellular responses and the longer-term evolutionary impacts of genome merging. In our study of newly formed triploid *C. reinhardtii* strains, RNA-Seq, flow cytometry, and qPCR results revealed significant transcriptomic and potential proteomic shocks, accompanied by disruptions in cytonuclear stoichiometry. Future studies focusing on the genomic changes occurring within these triploid lines will shed light into the potential molecular mechanisms, such as structural variation, genome fractionation, chromosomal instability and epigenetic modifications, providing deeper insights into the consequences of polyploidy.

## Acknowledgement

The authors thank Dr. Eylem Aydogdu for helping set up the Chlamydomonas system. We also thank Dr. Marlies Peeters and Dr. Zhen Li for their insightful discussions. YVdP acknowledges funding from the European Research Council (ERC) under the European Union’s Horizon 2020 research and innovation program (Grant No. 833522). YVdP and ODC received funding from the Fonds Wetenschappelijk Onderzoek (FWO Research Project – G0C0116N) and infrastructure grant (EMBRC Belgium, FWO ESFRI - I001621N). AVdV and FA-S were funded by Ghent University (Methusalem funding, BOF.MET.2021.0005.01). LP-B and QB were awarded PhD scholarships by the Fonds Wetenschappelijk Onderzoek (FWO) of Flanders (Grant Nos. 11H0426N and 1168420N, respectively).

## Author contributions

**Lucas Prost-Boxoen:** Conceptualization (lead), Formal analysis (lead), Investigation (equal), Writing - Original Draft (lead), Writing - review & editing (equal), Visualization (lead). **Quinten Bafort:** Conceptualization (supporting), Writing - review & editing (equal). **Antoine Van de Vloet:** Investigation (supporting), Formal analysis (supporting), Writing - Original Draft (supporting), Writing - review & editing (equal). **Fabricio Almeida-Silva:** Formal analysis (supporting), Visualization (supporting), Writing - Original Draft (supporting), Writing - review & editing (equal). **Yunn Thet Paing:** Investigation (equal). **Griet Casteleyn:** Investigation (equal). **Sofie D’hondt:** Investigation (equal). **Olivier de Clerk:** Conceptualization (supporting), Supervision (equal), Writing - review & editing (equal). **Yves Van de Peer:** Conceptualization (supporting), Supervision (equal), Funding acquisition (lead), Writing - review & editing (equal)

## Notes

### Competing Interest Statement

The authors have declared no competing interest.

