## Supplementary material for "Asymmetric genome merging leads to gene expression novelty through nucleo-cytoplasmic disruptions and transcriptomic shock in *Chlamydomonas* triploids": Supplmentary figure 1

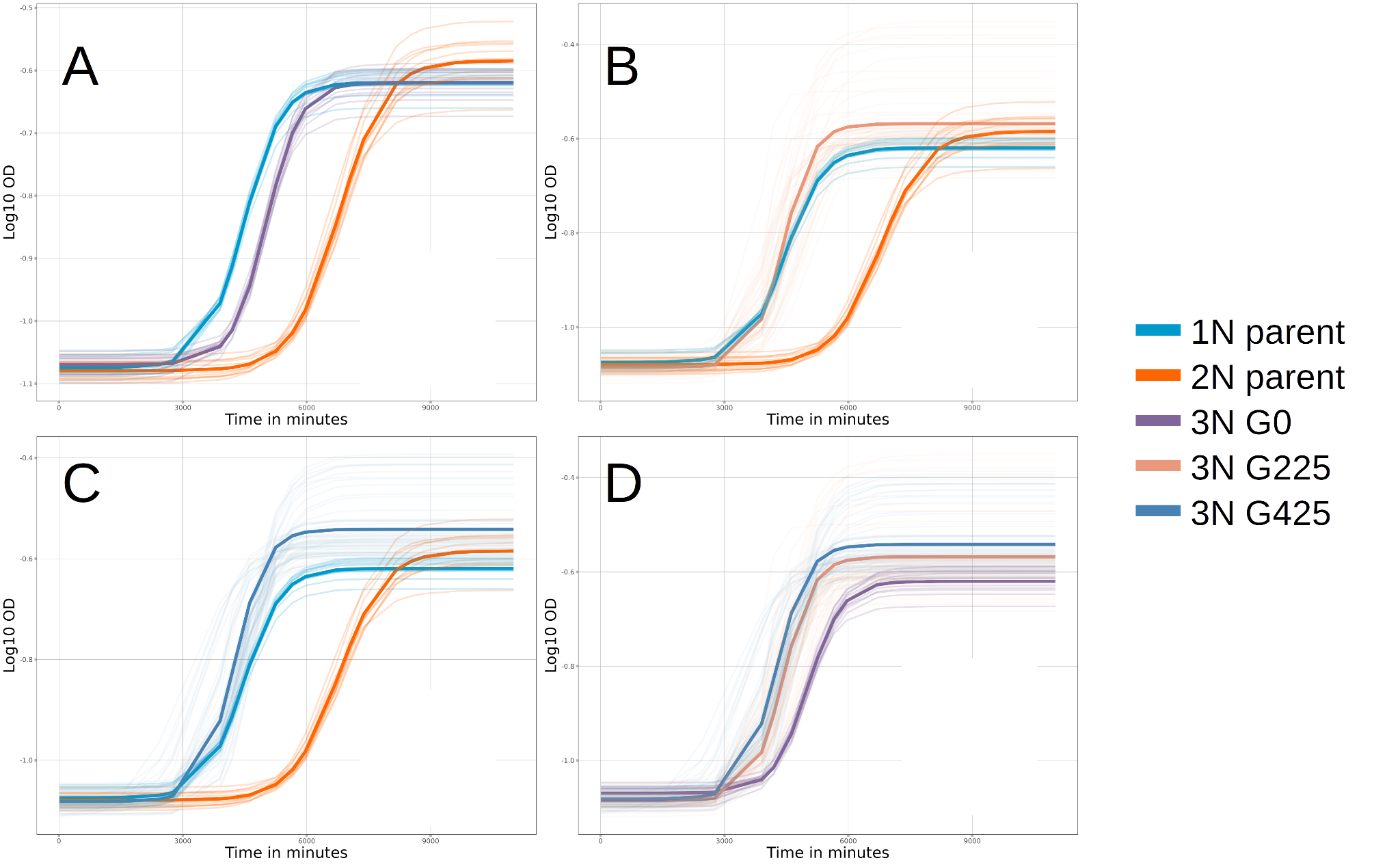

**Supplementary Figure S1.** Growth curves of the different experimental strains. This plot represent the growth curves of the haploid parent (1N), the diploid parent (2N), the triploid progeny (3N G0) and the triploid lines at generations 225 and 425 (3N G225 and 3N G425 respectively). Transparent lines represent the individual replicates, while bold lines represent the average growth curve for each strain. The growth curves are displayed in four separate plots for clarity and ease of comparison. **A**: growth curves of the two parental strains (1N and 2N) and the triploid progeny (3N G0). **B**: growth curves of the two parental strains (1N and 2N) and the triploid lines at generation 225 (3N G225). **C**: growth curves of the two parental strains (1N and 2N) and the triploid lines at generation 425 (3N G425). **D**: growth curves of the triploid lines at three different time-points, G0, G225 and G425.
