## Supplementary material for "Asymmetric genome merging leads to gene expression novelty through nucleo-cytoplasmic disruptions and transcriptomic shock in *Chlamydomonas* triploids": Supplmentary figure 2

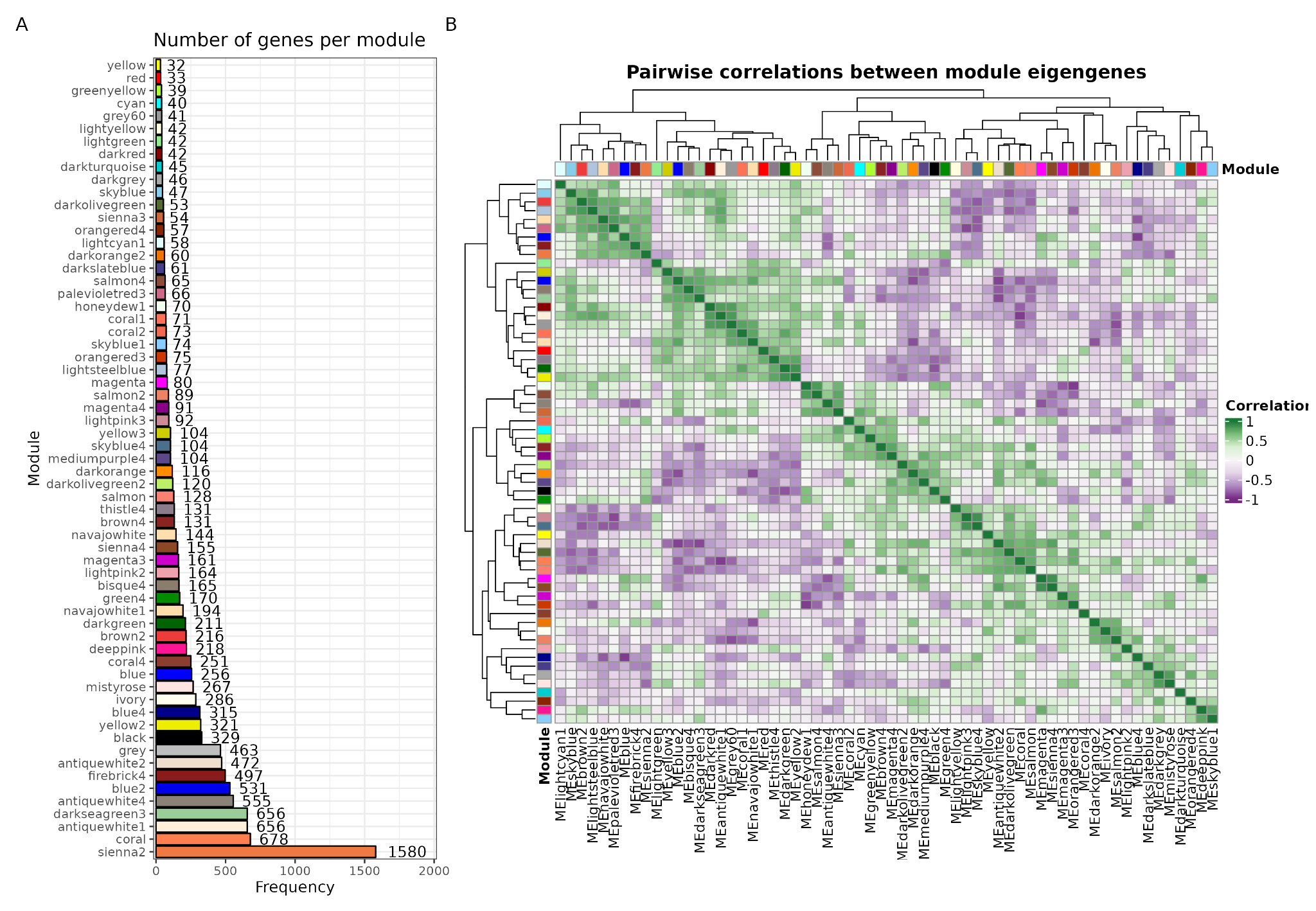

**Supplementary Figure S2.** Summary statistics of coexpression modules for the network with all samples. **A.** Absolute frequency of genes per module. **B.** Pairwise Spearman’s correlations between module eigengenes. Module eigengenes are the first principal component of each module, and they represent a summary of the expression profiles of the entire module.
