## Supplementary material for "Asymmetric genome merging leads to gene expression novelty through nucleo-cytoplasmic disruptions and transcriptomic shock in *Chlamydomonas* triploids": Supplmentary table 1

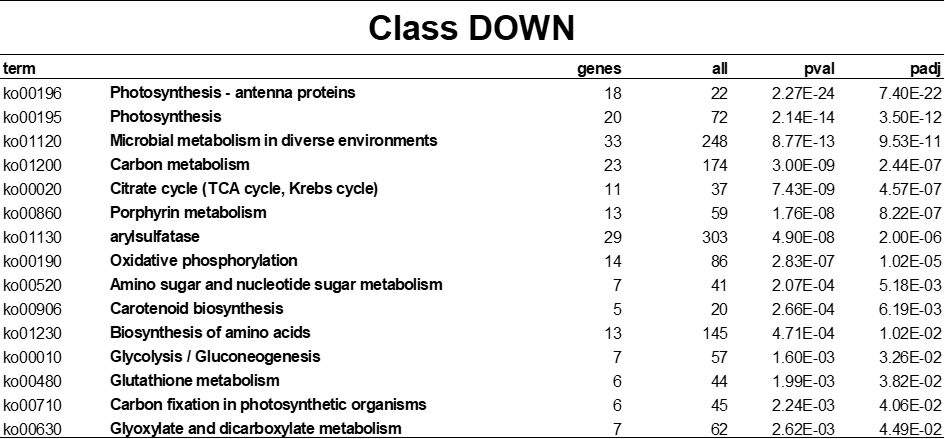

**Supplementary Table 1. Overrepresented KEGG pathways among persistent downregulated genes in the triploid.** This table lists KEGG pathways that are significantly overrepresented within genes exhibiting persistent decreased expression levels, organized in ascending order of adjusted p-values to highlight the most statistically significant enrichments.
