## Supplementary material for "Asymmetric genome merging leads to gene expression novelty through nucleo-cytoplasmic disruptions and transcriptomic shock in *Chlamydomonas* triploids": Supplmentary table 2

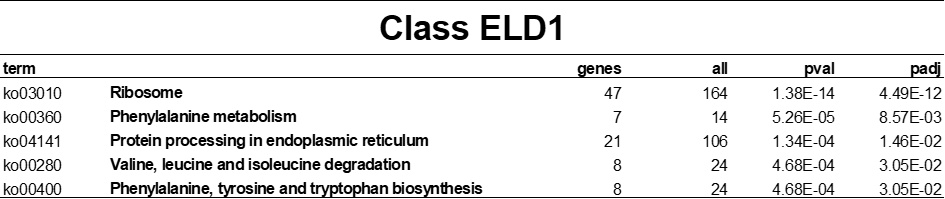

**Supplementary Table 2. Overrepresented KEGG pathways among genes showing persistent expression level dominance towards the haploid parent (ELD1).** This table lists KEGG pathways that are significantly overrepresented within persistent ELD1 genes, organized in ascending order of adjusted p-values to highlight the most statistically significant enrichments.
