## Supplementary material for "Asymmetric genome merging leads to gene expression novelty through nucleo-cytoplasmic disruptions and transcriptomic shock in *Chlamydomonas* triploids": Supplmentary table 3

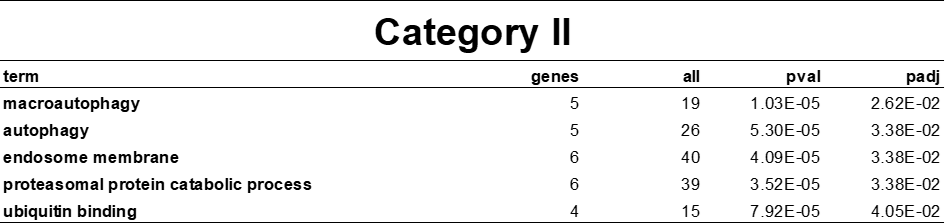

**Supplementary Table 3. Overrepresented GO terms in Category II genes.** This table presents Gene Ontology (GO) terms that are significantly overrepresented among Category II genes–those exhibiting a rapid increase in expression during the laboratory natural selection experiment. The terms are listed in ascending order of adjusted p-values to highlight the most statistically significant enrichments.
