## Supplementary material for "Asymmetric genome merging leads to gene expression novelty through nucleo-cytoplasmic disruptions and transcriptomic shock in *Chlamydomonas* triploids": Supplmentary table 4

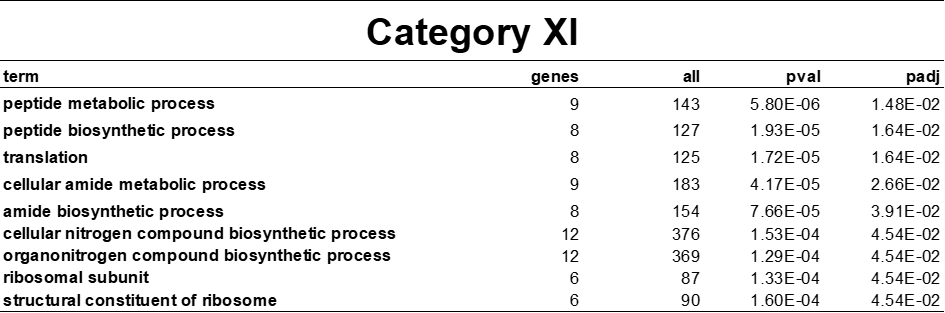

**Supplementary Table 4. Overrepresented GO terms in Category XI genes.** This table presents Gene Ontology (GO) terms that are significantly overrepresented among Category XI genes–those exhibiting a rapid decrease in expression during the laboratory natural selection experiment. The terms are listed in ascending order of adjusted p-values to highlight the most statistically significant enrichments.
